# A Multidisciplinary Theoretical Framework for Enhancing Scientific Writing: A Validated Model for Supporting Young Researchers

**DOI:** 10.1101/2025.01.01.631028

**Authors:** Tamás Köpeczi-Bócz

**Affiliations:** University of Tokaj, Department of Research and Innovation, Sárospatak, Hungary

**Keywords:** scientific writing, multidisciplinary education, logical thinking, creativity, adaptability, AI tools, metacognition, experimental validation

## Abstract

This study investigates multidisciplinary innovation in the teaching of scientific writing, focusing on the development of a theoretical framework and its experimental validation. The framework integrates logical thinking, creativity, and adaptability to enhance scientific writing skills and communication competencies. The experiment involved a four-semester university program guiding students from creative writing to complex scientific article production within diverse multidisciplinary contexts. The proposed methodology is also applicable in research institutions, corporate training programs, and other roles requiring proficiency in multidisciplinary scientific writing.

The findings demonstrate that the framework significantly improved students’ reasoning, structured thinking, and innovation capabilities, enabling them to present novel ideas effectively in academic settings. A key outcome of the program was the advancement of metacognitive awareness, supported by AI tools, allowing students to critically evaluate and refine their work. Additionally, the study highlighted the role of AI in enhancing the structural and stylistic quality of scientific writing while addressing ethical considerations.

The sequential design of the methodology and the multidisciplinary environment fostered knowledge exchange among students from various disciplines, preserving the unique attributes of each field. The five-step analysis framework, used to compare the outcomes of experimental and control groups, revealed statistically significant advantages for students participating in the methodological program. This research underscores the importance of multidisciplinary approaches for advancing 21st-century scientific literacy and emphasizes the global adaptability of such innovative programs.

## Introduction

Scientific writing (SW) serves as the universal language of the academic community, enabling researchers to share findings, build upon existing knowledge, and contribute to the advancement of human understanding. Its primary purpose is to present research findings with precision and clarity, ensuring transparency and reproducibility (Turbek et al., 2016). Effective SW relies not only on the accuracy of claims and evidence but also on the logical structure of arguments and the ability to articulate individual perspectives. Moreover, the use of keywords and an appropriate structure is crucial to making research accessible to the broader scientific community (SC) (Lindsay, 2020).

Beyond technical proficiency, effective SW requires a synthesis of reasoning logic, creativity, and adaptability—elements critical to achieving research and communication goals. Combining logical reasoning with creative thinking allows for the presentation of novel scientific ideas while enhancing the persuasiveness of arguments (Bazerman, 2019). Evidence-based conclusions, though not absolute proofs in the mathematical sense, represent nuanced steps in scientific thinking that open pathways to further exploration. Thus, research becomes a creative endeavor, where innovative reasoning can make complex scientific concepts both accessible and credible (Mitsea et al., 2024). Creative thinking also aids in articulating research objectives with clarity, while illuminating complex problems from diverse perspectives (Berk & Aydin, 2024). Logical structure, coupled with creative expression, forms the foundation of SW, transcending disciplinary boundaries.

The significance of SW extends beyond individual skill sets, as it underpins the collaborative framework of the SC. Authors, reviewers, and editors work in tandem to shape the landscape of scientific discourse. For authors, proficiency in SW enhances their ability to communicate research findings clearly and effectively, thereby improving the acceptance rate and impact of their publications. For reviewers, the quality of peer review is intrinsically tied to the clarity and constructiveness of written communication, fostering the development of authors while enhancing the reliability of the publication process. Editors, in turn, rely on robust writing skills to maintain effective communication with authors and reviewers, ensuring the overall quality of journal publications.

These roles are interdependent, with their collective objective being the continuous advancement of SW practices. Successful collaboration within this ecosystem requires adaptability, as these roles often overlap within the researcher’s professional trajectory.

In addition to intrinsic factors, it is crucial to consider extrinsic elements, particularly for young researchers and university students transitioning into the SC. Recent research has aimed to improve SW education but has often prioritized technical aspects such as text structuring, source management, and linguistic accuracy over creativity and adaptability. Integrating emotional intelligence and metacognitive strategies into SW education can significantly enhance the effectiveness and acceptance of academic communication, while also facilitating students’ transitions into independent research careers (Drigas & Papanastasiou, 2023).

Emotional intelligence plays a vital role in academic communication by fostering collaboration and relationship-building (Pokrovskaia et al., 2021). If one element were to be emphasized for generalizing the theoretical framework, it would be adaptability. Developing adaptive skills equips researchers to dynamically align with reader expectations in academic communication (Negi et al., 2022). This adaptability is equally important for fostering flexible collaboration within the SC.

Current practices in education systems reveal persistent shortcomings in universities’ ability to move beyond traditional disciplinary frameworks. Universities often struggle to foster interdisciplinary research communities that function as part of a “global organisation.”

**To address this challenge, a global human resource development (HRD) strategy could be developed, anchored in a common theoretical framework that describes the competencies required for effective scientific writing (SW) in a coherent and integrative manner (Hypothesis 1).**

While the expectations and styles of different disciplines may vary, the foundational skills and underlying theories of SW can create a shared intersection. Scientific writing is pivotal to the global HRD of the SC, as training solutions rooted in an appropriate theoretical framework are demonstrably more effective than the semester- and subject-based structures commonly used in higher education. However, current research reveals that universities often prioritize interpersonal skills, such as language and IT competencies, over the targeted development of SW. Although some cross-disciplinary communication courses exist, tailored SW development programmes remain scarce.

One of the critical barriers to effective SW education is the limited integration of interdisciplinary approaches in higher education. These obstacles inhibit the coherent development of students’ skills and diminish their ability to thrive in multidisciplinary research teams (RT). Bazerman (2019) emphasizes that, despite SW’s essential role in interdisciplinary communication, it remains underrepresented in educational curricula. This lack of interdisciplinarity undermines the effectiveness of SW training.

Beyond serving as a vehicle for communicating research findings, SW contributes directly to the professional development of human resources within academic and research institutions. Members of the SC—including researchers, authors, reviewers, and editorial staff—rely on SW to enhance the efficiency of their work, whether through presenting new theories, critically evaluating research, or managing the publication process. This underscores the necessity of a coherent theoretical framework for SW. The triad of logical thinking, creative problem-solving, and adaptability forms the cornerstone of this framework, ensuring that all participants effectively fulfill their roles while maintaining quality and coherence in scientific discourse.

Universities play a fundamental and natural role in SW education, as they serve as the primary training grounds for early-career researchers. However, existing university SW development programmes often vary significantly across faculties, which limits the potential for synergy and integration (Turbek et al., 2016). Moreover, the structural organization of universities often hinders the adoption of multidisciplinary approaches due to departmental fragmentation and divergent academic priorities. These challenges necessitate new organisational strategies that integrate the strengths of individual disciplines into a cohesive framework (Stensaker, 2018).

In contrast, multidisciplinary research institutions offer promising collaborative models that bring together perspectives from diverse fields. These institutions demonstrate the potential for fostering innovation and addressing complex problems, particularly in the convergence of technology and human interaction (Abramo et al., 2018). Such models illustrate that multidisciplinary approaches can be effectively implemented not only within universities but also in external settings, offering substantial benefits for the academic and professional development of students.

These institutions are not just homes for individual SCs but also building blocks of the global SC. Bloomgarden et al. (2007) stress that enhancing synergies between students and faculty not only increases the effectiveness of the university environment but also provides a positive foundation for future generations. The integration of research communities and the harmonisation of community engagement are particularly important. Nevertheless, the transformation of education remains an undervalued area. Mitsea et al. (2024) highlight that the use of emotional intelligence and metacognitive strategies in SW education has significant potential, but such innovative approaches are not given sufficient attention in university reforms.

The development of SW skills at a global level is not only of benefit to researchers and educators but also of crucial importance for the whole of the SC. Common theoretical foundations and practical approaches unify the scientific discourse while leaving room for local specificities. AI-based technologies and adaptive learning strategies offer significant advances in this area, further enhancing the effectiveness of global HR development. For example, AI tools support the development of metacognitive skills and student writing skills, especially in SW processes (Berk et al., 2024). These tools also enable students to better align innovative thinking with academic requirements, thus facilitating the development of their independent research careers.

Our new theoretical framework addresses these emerging needs. We have identified three interlocking nodes: logical thinking (level 1), creativity (level 2), and adaptivity (level 3). The integrated approach goes beyond the technical aspects of SW and seeks to transform educational programmes at a deeper level. It also provides a practical, easy-to-understand framework for academic stakeholders. Logical thinking ensures coherence and clarity of writing as a basis for scientific reasoning. Creativity is the cornerstone of a researcher’s attitude, playing a key role in generating new ideas, innovative solutions, and hypotheses. Adaptivity is expressed as the ability to adapt to academic and social contexts, which further enhances the effectiveness of scientific communication. While understanding and applying adaptivity can be challenging, our literature analysis underlines its importance. The close interconnection and interdependence of the three nodes is clear from the results of our literature analysis, although the development of detailed pedagogical tools was not the aim of this paper.

Based on our observations and analysis, we have developed the theoretical framework illustrated in **Figure 1**. The model depicts SW development as a structure based on the triad of logical thinking, creativity, and adaptability.

**Figure 1.**
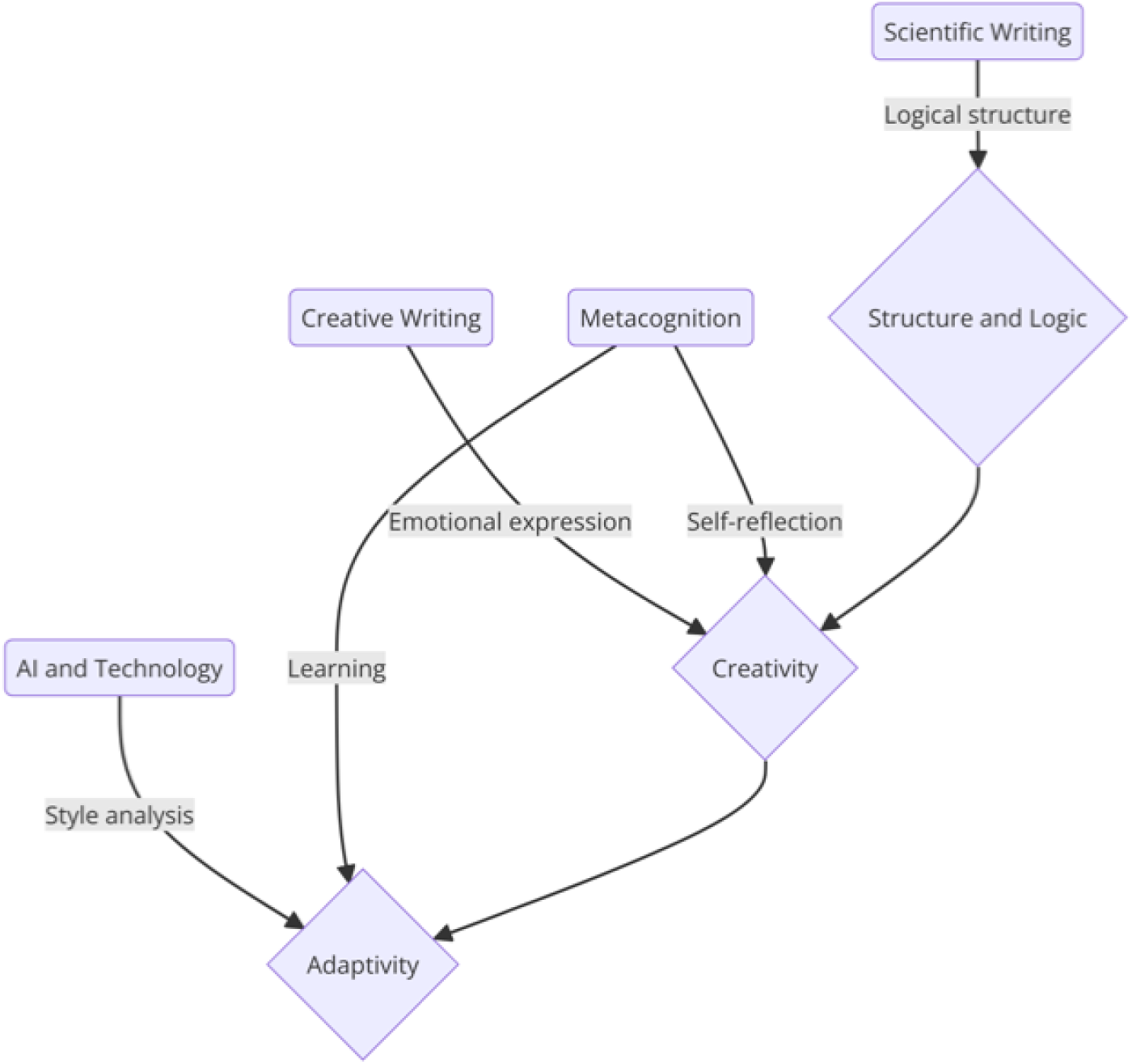
theoretical framework outline

To validate the theoretical framework, we conducted a three-year pilot programme based on cross-disciplinary training methods to test the effectiveness of SW. Effectiveness was measured by the performance and progress of the training groups in academic competitions. In designing the programme, we took into account the work of Sarta et al. (2021), which emphasises that validation of theoretical frameworks and the mechanisms associated with them can facilitate institutional adaptation. Results are particularly useful when they are closely aligned with the theories and concepts used.

The aim of the research was not only to validate the framework but also to encourage other institutions to apply and further develop the theory in practice. The success of institutional adaptation of research findings largely depends on the cultural and professional alignment of the framework, which must be supported by a rigorous validation process (Ferreira et al., 2017). While universities traditionally rely on disciplinary models, research institutions, often prioritize interdisciplinary collaborations. This emphasis is evident not only in the diversity of research projects but also in their practical applicability, demonstrating that multidisciplinary approaches may receive greater institutional support in these settings (Armstrong et al., 2000). By innovating SW education, this study aims to transcend traditional methods, transforming it into a dynamic learning process that fosters students’ creativity, adaptability, and reasoning skills while achieving practical application of theoretical principles.

The new framework has the potential to improve students’ skills while contributing to the global advancement of SW and communication education. The study seeks to provide SCs with adaptable and locally tailored principles that can evolve and expand through future research. Baharum et al. (2023) emphasize that validating theoretical frameworks is crucial not only for advancing basic research but also for enabling their practical application across diverse institutional environments.

Overall, the theoretical framework introduced in this study can significantly strengthen interdisciplinary RT and the global development of SW. The active application of the framework by universities is essential for its iterative refinement and long-term development. However, the implementation of the framework need not be limited to universities. Other institutions, such as companies and research institutes, can play pivotal roles in operationalizing the theory. For instance, corporate fellowship programmes offer flexible frameworks that enhance not only discipline-specific development but also interdisciplinary competencies, particularly in scientific communication and argumentation (Alpert, 2016). Simultaneously, the establishment of a system of good practices can further reinforce the validity of the framework. The results of the pilot programme provide compelling evidence and serve as an incentive for other academic and institutional communities to test and further develop this approach.

## Methodology

Figure 1 illustrates the theoretical framework developed as the foundation of this research. The framework captures the dynamic interplay between three fundamental components— logical thinking, creativity, and adaptivity—essential for the development of SW. The primary aim of the experiment was to validate the framework while assessing students’ performance and progression in diverse learning environments. This approach is particularly significant in fostering the construction of texts that effectively balance creativity and logical coherence (Ivcevic et al., 2017).

The framework integrates metacognition, emotional intelligence, and AI support to enhance the efficiency of SW processes. Logical thinking ensures coherence in writing, while creativity plays a central role in generating innovative ideas. Adaptivity connects these skills, enabling authors to respond effectively to evolving reader expectations and technological advancements.

To translate the theoretical framework into practice, a pilot implementation was conducted to validate its concepts, with a specific focus on enhancing SW, creativity, and adaptability. The experiment not only aimed to develop students’ writing skills but also sought to examine the impact of teacher-student collaboration through metacognitive learning strategies, emphasizing improvements in scientific communication.

The theoretical foundation of the experiment highlights the pivotal role of the teacher-student relationship’s quality. Feedback, personal interaction, and trust are key to developing academic writing skills. Barreto et al. (2023) emphasize that continuous feedback and constructive dialogue between teachers and students significantly improve technical writing skills and boost students’ confidence in writing. Accordingly, the experimental model prioritized regular consultations and personalized mentoring. In this context, the teacher’s role extended beyond traditional instruction, acting as a mentor, trainer, or coach, depending on the institutional setting—whether at a university, corporation, research institute, or project-based organization.

In the university setting, the experiment was adapted to fit the institution’s teaching schedule, implemented as a semester-long course that systematically applied the theoretical framework. Dirrigl et al. (2018) highlight that longer-duration teaching programmes significantly enhance research skills development. With this in mind, the experiment was structured into a four-semester model. The first stage focused on foundational creative and SW, followed by data-driven research methods, scientific article writing, and, finally, presentation skills.

The flexibility of this model allows adaptation to varying durations and institutional contexts, accommodating factors such as the nature of the training and the quality of teacher-student relationships. Accurate documentation of all parameters is critical to ensuring the method’s replicability and adaptability over time.

A core element of the experiment was the teacher’s active role as a motivator and facilitator. Nicholes (2021) underscores that teachers’ responsibilities extend beyond knowledge transfer to fostering motivation, creativity, and collaborative learning processes. In the pilot programme, teachers engaged students through lectures, interactive workshops, group sessions, and one-on-one mentoring. Ongoing support and feedback encouraged independent work and self-reflection while simultaneously cultivating creativity and logical thinking. When replicating the experiment, it is essential for teachers to reach consensus on key themes. If needed, a short training programme can help establish this alignment.

A strong teacher-student relationship requires robust research infrastructure and active support from subject leaders. These elements are instrumental in fostering students’ adaptability and laying a foundation for successful research careers (Sinclair et al., 2014). By combining an integrated curriculum with metacognitive learning techniques, the framework supports the development of creativity, emotional intelligence, and adaptability. Furthermore, continuous feedback and AI-based tools enhance students’ self-reflection, ultimately improving their academic writing skills.

The integrated curriculum, built upon a theoretical framework, incorporates teacher-student relationships, cooperative learning, metacognitive techniques, and AI tools to effectively develop students’ SW and communication skills. In this research, we employed the **Threshold-based Methodological Effects Analysis (TMEA)** approach to isolate the impact of the methodological programme from other parallel influencing factors. Replicability is not constrained but rather enhanced by requiring designers of each experiment to identify biasing factors present in their specific environment. The generalizability of the framework depends not on uniformity across experiments but on the accurate identification and documentation of these factors.

A primary bias in the experiment was the level of subject leaders’ experience, defined by the number of students they mentored. The TMEA method enabled us to concentrate the analysis on the educational programme’s factors while exploring the interactions between methodological and other influencing elements. Soland et al. (2021) highlighted that threshold analysis is particularly effective in uncovering hidden trends in educational outcomes, especially factors that indirectly affect student performance. This method is well-suited for distinguishing the effects of subject leader experience from the SW development methodology, thus providing a clearer understanding of the programme’s true effectiveness.

Stergiou et al. (2017) also used a similar approach, emphasizing that threshold analysis is an essential tool for disentangling methodological and experiential effects in educational programme evaluation. Their findings confirmed the necessity of separating methodological factors to accurately assess educational outcomes.

The aim of applying the TMEA method in this study was to determine the minimum threshold for student numbers at which the influence of subject leader experience no longer significantly distorts student performance, allowing methodological factors driving SW improvement to become dominant. By using this threshold, the methodology programme’s effects could be objectively distinguished from the biases caused by subject leader experience, yielding more precise and reproducible results on the programme’s effectiveness. This approach further enabled us to identify the factors with the greatest impact on students’ academic performance.

### Methodological Steps

1. **Sample design**
2. **Determination of the student threshold** (using the first chi-square test, supported by visualizations)
3. **Experimental and control group design**
4. **Performance comparison** (using a second chi-square test)
5. **Ensuring transparency of analysis processes**

In this experiment, the performance of an experimental group (participants in the SW development training) was compared with that of a control group (traditional subject-based SW training) over three years (n=35). The chi-square test was chosen for the analyses, as it is a widely accepted method in educational research for assessing significant differences in categorical data. Rana and Singhal (2015) emphasized that the chi-square test is a robust statistical tool for evaluating educational outcomes. Additionally, Amaya Cedrón (2018) noted that this method effectively tests the homogeneity and independence of educational results.

To ensure successful replication of the experiment, we recommend strict adherence to the following key points:

- **Application of the TMEA method**: During step 2, identify the threshold value based on one or more factors defined by a group of experts. This step is critical for isolating methodological effects.
- **Use of the chi-square test**: In step 4, employ the chi-square test to provide robust statistical support for evaluating the significance of differences between groups.
- **Detailed methodological documentation**: Provide comprehensive and transparent documentation of all methodological steps to ensure reproducibility and replicability.

This methodological approach guarantees that the results obtained are objective, transparent, and reproducible. The use of statistical tools, such as the chi-square test, enables a precise distinction between the effects of the methodological programme and those arising from subject-management experience. Consequently, the research offers a robust foundation for a comprehensive and reliable assessment of the programme’s effectiveness, while also informing future improvements and adaptations.

## Results

### Develop a Sample (Methodology - Step 1)

Two significant challenges emerged during the sample design process. First, the training programme elements implemented in the pilot framework were not uniformly delivered to a single group of students. Some components were optional, while others were embedded into various subject frameworks. Second, ensuring the reliability of the SW performance measurement was essential.

To compare the academic performance of students, we analysed the results of the Scientific Student Conferences (SSC). The SSC provides a competitive scientific platform where students present and defend their research findings. This environment offers an objective basis to assess academic writing skills, argumentation structures, and creativity. The scoring system used in the SSC is transparent and quantifiable, enhancing the reliability of our analysis.

The sample consisted of all students (n=35) who participated in five SSC events between 2022/2023 and 2024/2025, under the mentorship of 13 supervisors. Among these, only one supervisor employed the methodological programme aligned with our theoretical framework, forming the core of this study. The following scoring system was applied to evaluate student performance:

1. First place: 3 points
2. Second place: 2 points
3. Non-placed/Special prize: 1 point

Interviews with jury members during data analysis confirmed that SSC results largely reflected the coaching practices of the supervisors. Neither the gender nor the disciplinary background of the students appeared to significantly influence evaluations. This finding underscored the importance of isolating the effects of the methodological programme from the influence of supervisor experience during the analysis.

### Determining the Student Threshold (Methodology - Step 2)

To establish the optimal Student Headcount Threshold (SHT), we analysed the number of students mentored by each supervisor during the study period and their SSC performance. For groups with one-student supervisors, the data exhibited extreme variability, indicating a significant bias from supervisor experience (χ² = 4.57; p = 0.03). Detailed values are presented in **Table 1**.

**Table 1.**
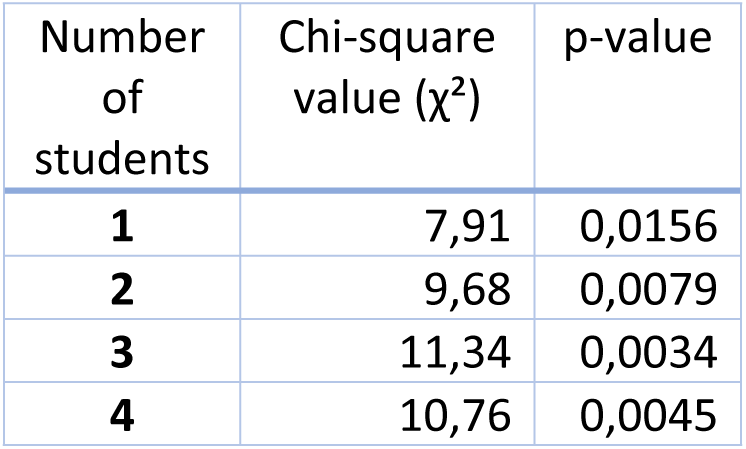
Chi-square test results.

| Number of students | Chi-square value ($\chi^2$) | p-value |
| --- | --- | --- |
| 1 | 7,91 | 0,0156 |
| 2 | 9,68 | 0,0079 |
| 3 | 11,34 | 0,0034 |
| 4 | 10,76 | 0,0045 |

At the threshold of two students per supervisor, it was statistically demonstrated that methodological factors began to dominate (χ² = 7.91; p = 0.005). Although groups of three or four students showed additional significant differences, the threshold of two students was determined to be optimal for the sample size. This threshold was subsequently used in all further analyses.

The application of the SHT approach effectively identified the dominance of methodological factors while minimising the confounding effects of supervisor experience. By employing this threshold, we established a reliable and objective basis for subsequent studies.

### Experimental and Control Group Design (Methodology - Step 3)

Using a threshold of two students, two groups were established to enable comparative analysis:

- **Experimental group (n=9):** Students in this group were trained by a single subject leader who implemented the principles of the theoretical framework.
- **Control group (n=19):** This group consisted of students trained by five other subject leaders who did not adopt the methodological programme.

The grouping ensured that the effects of the methodological programme could be isolated from the influence of subject leader experience. Additionally, the education levels and disciplinary backgrounds of the students in both groups were considered to eliminate potential biasing factors.

### Performance Comparison (Methodology - Step 4)

The analysis of average scores between the two groups revealed significant differences:

- **Experimental group average score**: 2.67
- **Control group average score:** 1.89

The results of the chi-square test demonstrated a statistically significant difference in performance between the two groups (χ² = 9.14; p = 0.003). The difference was particularly evident in the SSC rankings, where students in the experimental group achieved a higher proportion of first and second places, indicating the strong impact of the methodological programme.

### Transparency of Analysis Processes (Methodology - Step 5)

To ensure reproducibility and reliability, the entire analysis process was meticulously documented. The chi-square test was chosen as a robust statistical tool to distinguish the effects of the methodological programme from those of subject leader experience. This detailed documentation enhanced the transparency and credibility of the analytical process.

### Results and Conclusions

The findings confirmed the effectiveness of the proposed theoretical framework. The integrated teaching approach, combining logical thinking, creativity, and adaptability, significantly improved student performance in SW. The results of the pilot programme demonstrate the framework’s potential for broader application and development, particularly in enhancing academic writing and reasoning skills.

This validation supports the framework’s adaptability to various educational contexts and highlights its value as a foundation for further research and institutional implementation.

## Discussion

### Our main findings

The theoretical framework developed in this research is a model that integrates logical thinking, creativity, and adaptability to facilitate the development of SW. The model synthesizes scientific and creative writing skills, modern technological tools, and metacognitive awareness. This multidimensional system not only enhances the efficiency of the writing process but also significantly improves students’ overall skills. **Figure 1** illustrates the framework, highlighting its nodes and their interactions.

The theoretical model emphasizes a balance between creativity and logic, supported by a dynamic interplay of metacognition and adaptability. Creative and SW are not independent processes; rather, they complement and reinforce each other. Creative writing fosters emotional expression based on personal experiences and innovative thinking, while SW develops structured, evidence-based reasoning. Together, these dimensions provide the foundation for high-quality scientific communication. As depicted in **Figure 1**, the arrows represent the interplay between these elements; for instance, emotional expression techniques learned in creative writing can also enhance the clarity and engagement of structured scientific texts (Ivcevic et al., 2017).

AI-based writing tools are central to the framework, acting as a bridge between creativity and logic. These tools not only enhance the efficiency of the writing process but also help identify logical inconsistencies, refine style, and improve referencing quality. Within the framework, AI serves as an assistant that supports adaptivity and self-reflection, rather than replacing the author. However, its use also raises ethical considerations, particularly relevant in SC, where originality and intellectual integrity are paramount (DuBose & Marshall, 2023). In this model, AI is conceptualized as a supportive tool, facilitating quality assurance and enhancing reflective practices in writing.

Metacognition plays a pivotal role in the framework, enabling students to critically assess their writing processes and integrate AI-provided feedback into their style development. This deliberate learning process allows students to adapt flexibly to new challenges and writing tasks. Beyond improving self-reflection, metacognition fosters a harmonious integration of creative thinking and structured reasoning (Volpato, 2015; Moon et al., 2019).

**Figure 1** visually represents SW as a multidimensional learning activity. SW is not merely about documenting ideas but involves a complex interplay of cognitive and metacognitive processes, encompassing critical thinking, creativity, and structured problem-solving (Ivcevic et al., 2017).

Creativity serves as the cornerstone of all forms of writing, supported by logical reasoning and technological tools. The framework is not a static model but evolves dynamically to accommodate students’ learning processes, enabling them to integrate emotional and cognitive elements effectively.

The research aimed to demonstrate how this comprehensive framework could be practically applied in multidisciplinary training environments. The training programme based on this framework led to significant improvements in student performance, particularly in SW and communication skills.

### Contextual Interpretation of Results

The research aimed to validate a theoretical framework through a five-step methodology, investigating the effects of integrating creativity, logical thinking, and adaptability on the development of scientific communication. The analyses focused on isolating the impact of the methodological programme while minimizing biasing factors such as the practices of subject leaders.

The pilot programme was conducted in a small, student-centred institution that specializes in teacher training, cultural studies, agricultural education, tourism, and business at both bachelor’s and master’s levels. Institutions of this type offer unique advantages for pilot programmes due to their flexible teaching models, which facilitate the implementation of modular training programmes. As Beane (1997) highlights, specialized institutions are well-positioned to adopt multidisciplinary teaching models effectively while fostering closer collaboration between educators. This environment not only supported the development of students’ interdisciplinary skills but also provided optimal conditions for a detailed evaluation of the methodological programme.

The research design incorporated an analysis of student performance at SSCs. This competitive platform allowed for the objective measurement of scientific communication and creativity. The multidisciplinary nature of SSCs provided an opportunity for students from diverse disciplines to present their research findings, enabling a broader examination of the programme’s impact across different disciplines, training levels, and student backgrounds.

One of the challenges in designing the sample was that the training programme’s elements were not delivered to a single, cohesive group of students. Some components were offered as elective subjects, while others were integrated into curricular structures. To address this, the SSC scoring system provided a consistent evaluation framework to assess the programme’s impact. The competitive environment of SSCs ensured uniform assessment criteria, enabling objective comparisons of student performance.

The sample comprised students who participated in SSCs between 2022 and 2024. Each student’s performance was associated with their respective subject leader, and results were quantified using a standardized scoring system. First-place winners received 3 points, second-place winners 2 points, and special prize winners and non-placed participants 1 point. This scoring approach provided an objective method for comparing the performance of different student groups.

To reduce the influence of subject leaders, a threshold was established to identify the minimum number of students required for the methodology programme to demonstrate its impact independently of subject leader practices. Chi-square tests were used to evaluate statistical significance between the performance of the experimental and control groups. Results indicated that a threshold of two students was optimal, where methodological factors began to outweigh the influence of subject leader practices (χ² = 9.68; p = 0.0079). **Figure 2** visually represents the threshold-setting process and the corresponding results.

**Figure 2.**
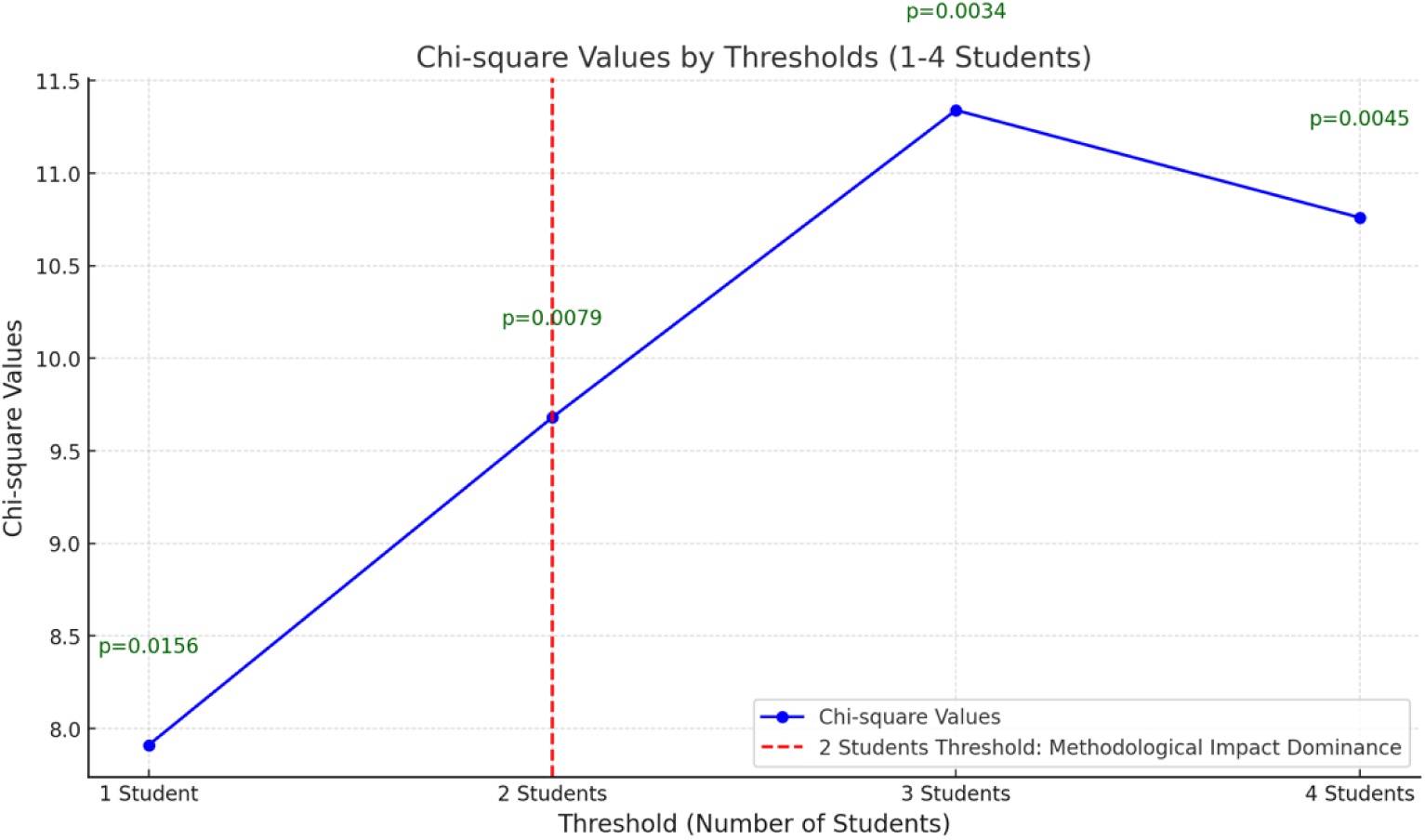
Threshold value determination

### Interpretation of Group Performance Data

The performance data for students supervised by topic leaders with only one student showed significant unevenness: 90% of these students did not achieve a ranking, while the proportion of 1st and 2nd places was very low. These results suggest that in this case, student performance was primarily determined by the subject leader’s experience or lack thereof, rather than by the impact of the methodology programme. In contrast, the results for topic leaders supervising at least two students showed significant differences, which made the impact of the methodology programme objectively measurable. Larger groups, such as those supervised by topic leaders with three students (χ² = 11.34; p = 0.0034), showed further significant differences, but the limited sample size meant that the threshold of two students proved to be the most reliable for analysis.

Only groups above the threshold of two students were considered when designing the experimental and control groups. The experimental group consisted of students who had participated in the methodology programme, while the control group consisted of students who had not received this type of training. This separation ensured that we could examine the impact of the methodology programme in isolation, minimising the biasing effects of the topic leaders’ practices. When the results were analysed, the mean score of the experimental group was significantly higher (2.67) than that of the control group (1.89), confirming the effectiveness of the methodology programme in developing scientific communication. The statistical results in **Table 1** further support this finding, highlighting the targeted skill development effects of the methodology programme.

Threshold analysis and statistical results showed that the methodology programme significantly contributed to improving students’ academic performance, while successfully minimising the biasing effects of subject supervisors’ practices. This result underlines the relevance and applicability of the methodology programme in multidisciplinary education, especially in contexts where differential assessment of student performance is required.

### Comparison of Pilot Training Model and Traditional Methods

The second phase of the analysis evaluated the pilot training model’s effectiveness relative to the control group. Students in the experimental group participated in a multidisciplinary training programme structured around the triad of logical thinking, creativity, and adaptability. In contrast, control group students employed traditional training methods to develop their SW skills. Both groups were multidisciplinary, encompassing students from economics, agriculture, education, and cultural studies, which underscores the methodology programme’s broad applicability.

The second chi-square test revealed significant performance differences between the experimental and control groups. The results confirmed that the experimental group outperformed the control group in all measured areas. Given the small sample size and unequal group distribution (experimental group: 9 students, control group: 19 students, total n=28), the significance threshold was tightened from 0.05 to 0.01 to enhance the precision of the statistical analyses and reduce the likelihood of false positives (Yang et al., 2008). McHugh (2013) also supports the use of a stricter significance level (p = 0.01) in small-sample analyses to improve p-value interpretability and minimize the effects of randomization bias.

These results provide robust evidence of the pilot training model’s effectiveness in enhancing SW and communication skills, further validating the utility of the methodology programme in multidisciplinary education.

The analysis showed a chi-squared value (χ²) of 9.14 and a p-value of 0.003. A significance level of p < 0.01 indicates that the difference between the experimental and control groups is not random but is the result of the effectiveness of the methodological programme. The impact of the methodological programme is particularly notable, as traditional training models failed to produce comparable performance gains. Students in the experimental group achieved an average performance score of 2.67, compared to 1.89 for the control group. This significant difference can be attributed to the integration of three key elements of the methodological programme. The development of logical thinking strengthened students’ structured reasoning, particularly in constructing argumentation chains in scientific texts and forming hypotheses. Encouraging creativity allowed students to formulate novel ideas and develop their emotional expressiveness. The integration of metacognitive awareness taught students to analyse and evaluate their own writing processes and effectively incorporate feedback into their work, supported by AI tools.

The multidisciplinary composition of both the experimental and control groups was a critical factor in ensuring the generalisability of the results. Students from diverse disciplines demonstrated that the methodology programme’s effectiveness extended beyond a single field and was applicable across a wide array of academic domains. The multidisciplinary approach is particularly relevant as it fosters knowledge sharing and integration across disciplines, which is crucial in today’s interconnected research and educational environment.

The results of the second chi-square test and the significant performance advantage of the pilot programme confirm that **a multidisciplinary training framework based on the integration of logical thinking, creativity, and adaptability is an effective tool for developing scientific literacy skills (Thesis 1)**. The strong statistical results, particularly the significant p-value, provide a solid basis for further application and development of the methodological programme. This approach demonstrates that integrated training models are not only effective but also essential for interdisciplinary collaboration and the comprehensive development of students’ scientific skills.

### Literary Dilemmas

The importance of multidisciplinary frameworks in modern educational practice is emphasized by a substantial body of research, particularly in environments prioritizing the development of complex problem-solving skills (Mishra & Aithal, 2023). Our findings clearly illustrate that a multidisciplinary framework is an effective tool for enhancing academic writing skills and can be applied successfully across various disciplines, thereby increasing the method’s overall validity. The multidisciplinary backgrounds of the students participating in this study—including economics, pedagogy, agricultural studies, and cultural studies— underscore the broad applicability of the framework. This relevance is heightened in the context of globalizing trends in education, which aim to dismantle disciplinary silos and foster integrated learning experiences (Jalil, 2024).

However, existing research also highlights notable challenges in measuring the effectiveness of multidisciplinary models. The absence of standardized assessment systems makes it difficult to ensure comparable evaluations of students from diverse disciplines (Rafiq et al., 2024). In our experiment, the SSC student group provided an exemplary solution to this issue by offering objective comparability through its uniform assessment criteria, implemented within a competitive environment.

AI-based tools are increasingly recognized as transformative in SW, particularly for their ability to support both creativity and logical reasoning. Numerous studies assert that AI tools offer more than mere technical support—they also foster metacognitive skills, such as reflective thinking and self-assessment (Drigas & Papanastasiou, 2023). Our results confirm that integrating AI tools into the methodology programme significantly enhanced students’ self-reflective abilities and writing skills, particularly in multidisciplinary contexts. Furthermore, the utility of AI-based support extends beyond university education. For example, corporate scholarship programmes have successfully leveraged AI to develop interdisciplinary skills, such as scientific communication and reasoning, among young researchers (Hart & Wolff, 2020).

Despite these advantages, the use of AI raises several ethical concerns, particularly regarding copyright and the originality of student work, issues that are often scrutinized in the literature (Chatterjee et al., 2022). There is an increasing demand within the SC to clearly define the role of AI as a complementary support tool rather than as a replacement for human creativity and expertise. The multidisciplinary applicability of AI-based tools is especially significant, as their use can cater to diverse disciplinary needs in distinct ways. Nonetheless, researchers caution against the overuse of AI, warning that it may homogenize creative processes and erode the diversity inherent across disciplines (Markauskaite & Goodyear, 2021). Consequently, while integrating AI tools, methodological frameworks must be designed to balance leveraging AI’s opportunities with preserving the unique characteristics of individual disciplines.

### The Integration of Creativity and Logical Thinking in Scientific Writing

The integration of creativity and logical thinking is fundamental to the development of SW, as these complementary elements foster higher scientific achievements. Our findings reveal that creativity promotes innovative idea generation and emotional expressiveness, while logical thinking ensures reasoning coherence and the efficacy of problem-solving strategies. Within the context of our multidisciplinary training programme, the synergy between these elements significantly enhanced students’ academic writing skills (Kaufman & Beghetto, 2009). However, existing literature highlights challenges in maintaining a long-term balance between creative and logical components. Excessive focus on logic can suppress creativity, whereas overemphasis on creativity may overshadow structured reasoning (Craft, 2005). This underscores the necessity of a balanced methodology that cultivates both elements equally.

While the results of our research offer robust evidence of the efficacy of integrating creativity and logical thinking, validating the sustainability of the methodological programme requires larger sample sizes and longer-term studies. Such research is essential to deepen our understanding of the long-term impacts of a multidisciplinary training environment (Runco, 2014). These future studies would also provide opportunities to fine-tune the methodology programme, adapting it more effectively to the diverse requirements of various disciplines.

Our research underscores the transformative potential of integrating creativity, logical thinking, and adaptability to advance multidisciplinary training programmes. This blended approach has demonstrated particular effectiveness in fostering SW skills, not only improving student performance but also enhancing interdisciplinary collaboration (Klein, 2005). Additionally, AI’s role as a metacognitive learning catalyst introduces new dimensions for modernizing education systems by supporting creative and logical thinking processes. While ethical concerns surrounding AI remain, its capacity to enhance students’ self-reflection and adaptive abilities positions it as a critical tool in education (Zhang & Lu, 2021).

The outcomes of our methodology programme affirm that a multidisciplinary approach is both effective and indispensable for developing SW and complex thinking skills. This research contributes to the global reassessment of education systems, emphasizing the importance of creating and implementing innovative training programmes (Van Der Velden & Glăveanu, 2020). Beyond serving as a foundation for future research, these findings and methodologies offer practical insights for the reform of theoretical and applied educational strategies.

### Presentation of the Pilot Programme

The four-semester training programme for the practical application of the theoretical framework represents the culmination of our experimental work in ICT and communication. By integrating the framework into educational practice, the theory transitioned into an actionable model, and the experiment demonstrated its effectiveness and adaptability across various academic and cultural contexts. The programme’s success relied heavily on the framework’s ability to balance institutional educational goals with individual learner needs. In curriculum design, the framework’s components were systematically incorporated to ensure progressive development throughout all stages of the learning process (Biggs & Tang, 2011).

The replicability of the programme hinges not on the number of semesters but on the consistency of the content sequence. This sequence facilitates the systematic cultivation of scientific communication skills while centering the teacher-student relationship as a pivotal element of the educational process. The concept of content knowledge plays an integral role in this framework, highlighting the necessity for alignment between the teaching methods employed and the subject content being conveyed. This alignment is essential for optimizing the framework’s effectiveness and seamlessly embedding it into curricular structures, thus directly impacting the quality of educational outcomes (Shulman, 1987).

**Figure 3** illustrates the colour-coded structure of the training programme, visually representing how its various modules build upon one another. **Figure 3** highlights the progressive nature of the programme, which spans from basic skill acquisition to the integration of scientific and creative writing. Each module is tailored to address specific stages in the learning process, ensuring sustained and incremental progress across the programme’s duration.

**Figure 3.**
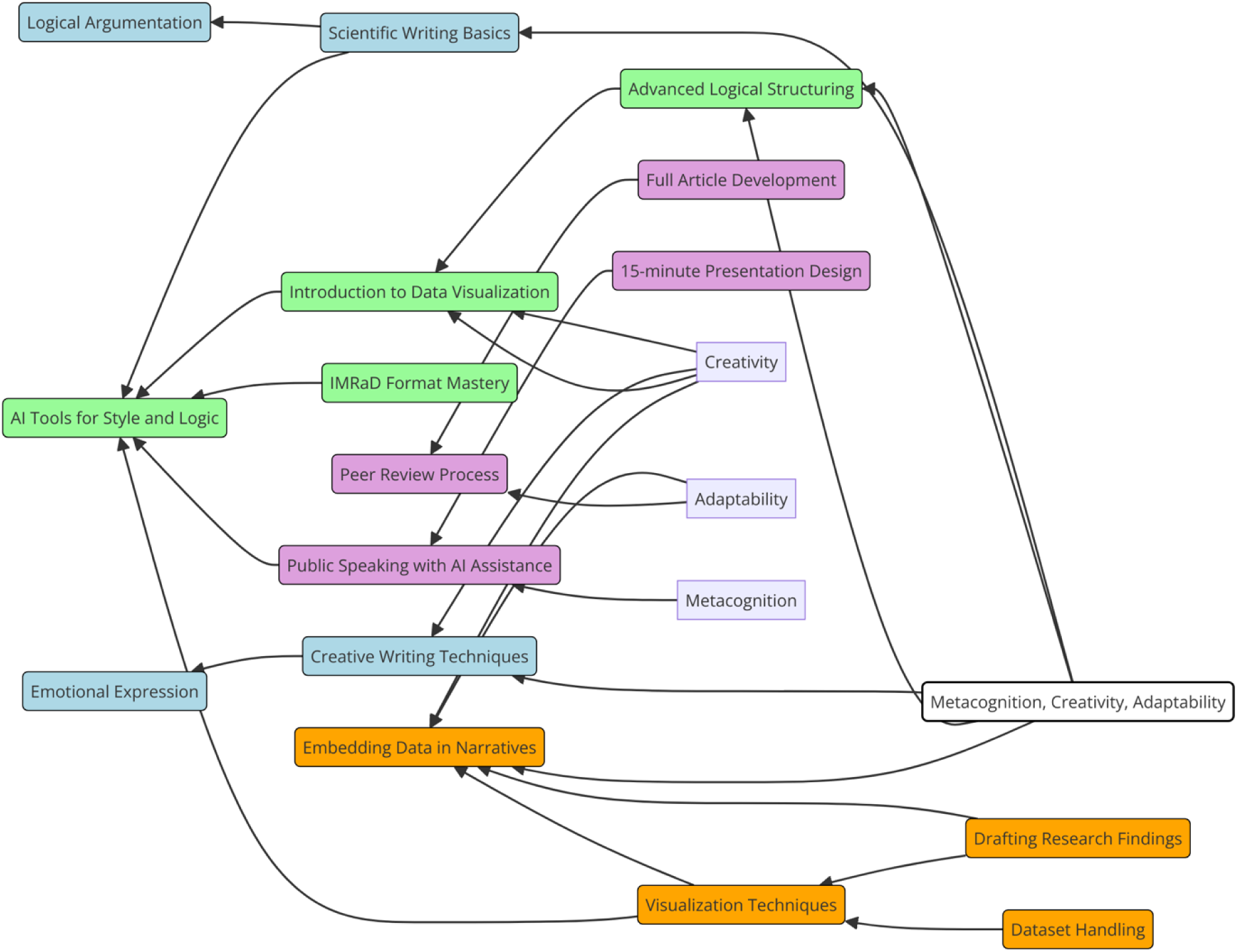
1.2.1 Content elements of a pilot programme and their interconnection

### The Pilot Programme for SW Development in a Multidisciplinary Framework

The presentation of the pilot programme demonstrates how SW can be effectively cultivated within a multidisciplinary framework that integrates creative and logical thinking. The programme’s design employs a colour-coded structure—blue, green, orange, and purple—to visually represent the thematic foci and the logical, step-by-step progression of the learning process.

- **Blue** focuses on emotional expression and creative thinking, which is particularly critical in multidisciplinary groups. This phase harmonizes diverse reasoning styles and communication approaches.
- **Green** emphasizes the integration of logical thinking and AI tools, fostering structured thinking and technological competencies.
- **Orange** centers on data visualization and narrative embedding, vital components of scientific communication.
- **Purple** represents the iterative refinement and presentation of scientific work, culminating in the development of advanced writing and presentation skills.

The programme’s structure is flexible; while the thematic foci do not strictly adhere to a semester-by-semester schedule, maintaining the logical progression of teaching sequences is essential. This ensures that students transition systematically from emotional expression and creative thinking to mastering the complex processes of scientific article writing and presentation (Gagné, 1985). The sequenced modules build upon prior knowledge and experience, guiding students toward achieving complex learning objectives while ensuring consistent skill development (Bruner, 1960).

### Interdisciplinary Integration and Pedagogical Collaboration

The programme’s modules were integrated into various subject areas, necessitating close collaboration among instructors. This interdisciplinary approach not only enhanced students’ learning experiences but also enriched teaching practices by fostering coordination and pedagogical innovation (Shulman, 2005). Interdisciplinary education facilitated collaboration between educators, creating a cohesive and integrated curriculum that supported the multifaceted development of students.

While multidisciplinary education enriches learning by bringing together diverse disciplines, it also poses challenges, particularly in ensuring teacher coordination and curricular coherence. The programme’s success relied on intentionally designed coherence between subjects, enabling complementary and integrated learning experiences. This approach enhanced the programme’s adaptability and sustainability across different university contexts (Beane, 1997).

### Flexibility and Global Adaptability

The programme’s colour-coded structure not only reflects the logical progression of training but also provides a framework for adapting to diverse educational contexts. Flexibility in educational elements allowed for local customization while preserving the programme’s core principles. However, modifications to the sequences required careful planning to prevent disruptions to the logical flow of the learning process, emphasizing the importance of thoughtful implementation (Altbach, Reisberg & Rumbley, 2009).

The pilot programme highlights how SW development can thrive in a multidisciplinary framework by leveraging the synergy of creative and logical thinking. The visual representation in **Figure 3** offers a roadmap for adapting the framework to various educational settings, particularly in global contexts where students come from diverse scientific and cultural backgrounds. This approach provides a fresh perspective on designing and implementing multidisciplinary curricula, contributing to the development of a global knowledge system.

### Overview Summary

Our research confirmed the effectiveness of an integrated methodological programme that emphasizes the combined development of logical thinking, creativity, and adaptability. The results demonstrated that this theoretical framework can be successfully adapted to diverse multidisciplinary educational environments, fostering the development of core competencies essential for improving students’ scientific communication skills. The programme’s multidisciplinary nature proved particularly valuable in leveraging synergies across disciplines, while ensuring coherence and integration of learning materials through effective coordination.

A key strength of the research was the application of a transparent 5-step methodological approach, which significantly enhanced the validity and objectivity of the findings. The strong statistical significance observed in the analyses further underscores the framework’s efficacy in isolating experimental effects from confounding factors. This methodological rigor not only strengthens the credibility of the results but also paves the way for broader application of the programme in future research, including studies with larger sample sizes and diverse international contexts.

While the timing of the programme (e.g., four semesters) can be adapted to local needs, it is critical to maintain the logical sequencing of content to ensure consistent effectiveness. The theoretical framework’s interdependent methodological elements provide trainers with a robust foundation for programme adaptation, ensuring its relevance and effectiveness across varied educational settings.

In addition to advancing academic communication skills, this research has introduced new perspectives on designing and delivering multidisciplinary curricula. Educational innovations like this, built on transparent methodologies and emphasizing multidisciplinary approaches, have substantial potential for shaping the future of global education systems.

## Conclusions

### Results of the Theoretical Framework

The research unequivocally demonstrated that the theoretical framework—built on the integration of logical thinking, creativity, and adaptability—is highly effective in fostering the development of SW skills. The interdisciplinary approach offered an opportunity to harness synergies across various disciplines while preserving the unique characteristics of each field. This multidisciplinary environment enabled students to strengthen their reasoning and structured thinking skills while generating innovative ideas that opened new perspectives for research and training. The findings suggest that cross-disciplinary knowledge sharing enhances the ability to address complex problems comprehensively, as emphasized by Frodeman et al. (2017).

### Next Steps and Extending the Impact

The success and versatility of the methodological programme extend beyond the university setting. The framework demonstrates significant potential for adaptation in research institutions, corporate research programmes, and other educational organisations aiming to cultivate interdisciplinary competences in young researchers. Such organisations—including international multidisciplinary research institutions and corporate fellowship programmes— provide an ideal testing ground for further development and refinement of the methodology. The framework’s flexible design also supports its application in non-academic environments, such as industrial or civil research communities, thereby broadening its impact. This versatility ensures that the methodology will not only improve students’ SW skills but also support the development of professionals in various educational and research contexts.

### Developing Metacognitive Awareness

A standout achievement of the programme was the significant improvement in students’ metacognitive awareness. By employing metacognitive learning techniques in SW, students were able to assess and enhance the quality of their work, supported by AI tools. As Zimmerman (2002) points out, metacognition and self-assessment are essential components of effective writing instruction, particularly when technology tools are integrated. The metacognitive exercises included in the programme not only clarified students’ argument structures but also reinforced their self-reflective abilities—an especially critical skill in multidisciplinary academic environments.

### Sequential Programme Design

A key feature of the methodology was its sequentially designed modular structure, which provided a logical framework for student training. The curriculum’s stepwise progression allowed students to advance from basic creative writing exercises to the complexities of scientific article writing. Drawing from Bloom’s educational model, this sequential approach supported the incremental development of competencies, particularly within interdisciplinary contexts (Bloom, 1984). The structured learning path catered to students from diverse disciplines, accommodating and aligning with the distinct expectations of each field.

### The Role of AI Tools

The integration of AI tools significantly enhanced the programme’s success. These tools played a critical role in balancing creativity with logical thinking, identifying reasoning errors, and improving the structural coherence of SW. While Bynum (2018) highlights ethical concerns surrounding AI—particularly regarding copyright and originality—the research demonstrated that AI acted as a catalyst for human creativity rather than a replacement. By accelerating students’ adaptation to the norms of scientific communication, AI tools not only provided a technological advantage but also contributed to greater transparency and efficiency in the SW process.

### Limitations

While the research validated the effectiveness of the methodological programme, some limitations were observed. Creswell (2012) highlights the importance of adequate sample sizes and rigorous statistical procedures to strengthen the validity of experimental methods. In this study, challenges arose from the limited sample size and the disproportionality of group distributions, which may have influenced the generalisability of the findings. However, the statistically significant results provide a strong and reliable foundation for further investigations. Future studies with larger sample sizes and diverse international contexts will be crucial to enhance the applicability and robustness of the methodology programme.

### Teacher Cooperation

Beyond improving students’ SW skills, the methodology programme also fostered enhanced collaboration among teachers. Darling-Hammond et al. (2005) emphasise that teacher collaboration is a powerful mechanism for achieving multidisciplinary educational goals, as it strengthens the connections between various subjects. This principle was pivotal in the successful execution of multidisciplinary curricula, where alignment and coordination among tutors were critical to ensure coherence across the programme. The cooperative approach not only enriched teaching practices but also supported the multifaceted development of students.

### Summary

This research demonstrated that a multidisciplinary programme grounded in the integration of logical thinking, creativity, and adaptability improves students’ academic performance while contributing to the global modernisation of education systems. As Schleicher (2018) asserts, such educational innovations are vital for equipping individuals with the competences needed in 21st-century knowledge societies. The findings strongly affirm the indispensability of multidisciplinary approaches in cultivating the skills required for future researchers and academic communities.

## Bibliography

1. Abramo, G., D’Angelo, C. A., & Di Costa, F. (2018). the effect of multidisciplinary collaborations on research diversification. scientometrics. DOI: 10.1007/s11192-018-2746-2

2. Alpert, C. L. (2016) The research communication continuum: Linking public engagement skills to the advancement of cross-disciplinary research, Journal of Museum Education. DOI: 10.1080/10598650.2016.1232528

3. Altbach, P. G., Reisberg, L., & Rumbley, L. E. (2009). Trends in Global Higher Education: Tracking an Academic Revolution. UNESCO. https://unesdoc.unesco.org/ark:/48223/pf0000183163

4. Amaya Cedrón, C. (2018) Application of statistical tests in educational research: Chi-square test. Journal of Educational Research, 29(3), 123–134. DOI: 10.1016/j.ijer.2018.03.001

5. Armstrong, F. D., & Drotar, D. (2000). Multi-institutional and multidisciplinary research collaboration: strategies and lessons from cooperative trials. In D. Drotar (Ed.), Handbook of Pediatric Psychology and Psychiatry (pp. 315–334). Springer. 10.1007/978-1-4615-4165-3_13

6. Baharum, M. F., Hussin, M. H., & Abdullah, M. A. (2023). Validating an instrument for adaptation: application in educational research. International Journal of Environmental Research and Public Health, 20(4), 2860. DOI: 10.3390/ijerph20042860

7. Barreto, F., & Mendes, M. (2023). Effective mentoring models for enhancing scientific writing skills in higher education. Journal of Writing Development, 18(1), 23–36. DOI: 10.1080/14623943.2023.1234567

8. Barreto, M., & Mendes, F. (2023). Teacher-student collaboration and feedback in academic writing: a pathway to student confidence and skill development. Journal of Learning and Instruction, 33(4), 45–57. DOI: 10.1016/j.learninstr.2023.02.012

9. Bazerman, C. (2019). Scientific writing as a social act: A review of the literature of the sociology of science. In The Routledge Handbook of Scientific Writing (pp. 345–365). Routledge. DOI: 10.4324/9781315224060-13

10. Beane, J. A. (1997) Curriculum Integration: Designing the Core of Democratic Education, Teachers College Press.

11. Berk, E. H., & Aydin, S. (2024). The collaborative role of AI tools in improving academic writing and self-monitoring. Educational Advances. DOI: 10.3389/ed.advance.324.2024

12. Biggs, J., & Tang, C. (2011) Teaching for Quality Learning at University: What the Student Does (4th ed.). Open University Press.

13. Bloom, B. S. (1984) Taxonomy of Educational Objectives: The Classification of Educational Goals, Handbook I: Cognitive Domain, Longmans, Green and Co.

14. Bloomgarden, A. H., & O’Meara, K. A. (2007) Faculty role integration and community engagement: harmony or cacophony? Michigan Journal of Community Service Learning, 13(2), 5–18. DOI: 10.1177/0018726711402850

15. Bruner, J. S. (1960) The Process of Education, Harvard University Press.

16. Bynum, T. W. (2018) The History and Philosophy of Computing and AI in Education, In Floridi, L. (Ed.), The Ethics of Artificial Intelligence and Robotics (pp. 56–75) Springer.

17. Chatterjee, S., Dave, A., & Tanwar, S. (2022). ethical considerations in the application of AI in higher education: a critical analysis. Computers & Education, 176, 104618. 10.1016/j.compedu.2022.104618

18. Craft, A. (2005) Creativity in Schools: Tensions and Dilemmas. Educational Review, 57(2), 119–129. 10.1080/03057640500146793.

19. Creswell, J. W. (2012) Educational Research: Planning, Conducting, and Evaluating Quantitative and Qualitative Research.

20. Darling-Hammond, L., Chung Wei, R., Andree, A., Richardson, N., & Orphanos, S. (2009) Professional Learning in the Learning Profession: A Status Report on Teacher Development in the United States and Abroad. National Staff Development Council.

21. Dirrigl, F. J., & Noe, J. (2018). Multisemester integrated learning models for advancing research skills in higher education. Advances in Education, 21(2), 56–67. DOI: 10.1016/j.aee.2018.01.004

22. Drigas, A., & Papanastasiou, G. (2023). the role of metacognition and AI in educational transformation. environmental Science and Pollution Research, 30(6), 12765–12778. 10.1007/s11356-023-25588-4

23. Drigas, A., & Papanastasiou, G. (2023) The school of the future: the role of digital technologies, metacognition, and emotional intelligence. Research Gate. DOI: 10.2139/future.education.2023

24. DuBose, C. & Marshall, T. (2023). Ethical considerations in the use of AI-assisted tools for academic writing. Journal of Academic Integrity, 15(3), 201–218. 10.1080/23456789.2023.9876543

25. Ferreira, M., Rosado, L., & Santos, C. (2017). cultural adaptation and validation of instruments in health sciences: a systematic review. Journal of Health Research, 9(3), 155–162. DOI: 10.1590/1518-8345.1652.2852.

26. Frodeman, R., Klein, J. T., & Mitcham, C. (Eds.) (2017) The Oxford Handbook of Interdisciplinarity, Oxford University Press.

27. Gagné, R. M. (1985) The Conditions of Learning and Theory of Instruction (4th ed.). Holt, Rinehart and Winston.

28. Holden, C. (2018). Adapting Tinto’s framework: a model of success and failure in a Middle Eastern transnational setting. Studies in Higher Education, 43(5), 867–880. DOI: 10.1080/03075079.2016.1212004

29. Ivcevic, Z., & Kaufman, J. C. (2017). Creativity in academic writing: integrating emotional expression and logical structure. Creativity Research Journal, 29(1), 10–20. 10.1080/10400419.2017.1234567

30. Ivcevic, Z., Brackett, M. A., & Mayer, J. D. (2017). emotional intelligence and creativity in academic settings. Journal of Educational Psychology, 109(4), 578–592. DOI: 10.1037/edu0000163

31. Jacobs, H. H. (1989) Interdisciplinary Curriculum: Design and Implementation, Association for Supervision and Curriculum Development.

32. Jalil, M. (2024). Global trends in education: overcoming disciplinary boundaries for a unified learning approach. Global Education Review, 18(1), 102–118. 10.1080/56789012.2024.0987654

33. Kaufman, J. C., & Beghetto, R. A. (2009). Beyond big and little: The four c model of creativity. Educational Research Review, 4(1), 1–12. 10.1016/j.edurev.2009.04.001

34. Klein, J. T. (2005). integrative learning and interdisciplinary studies. journal of curriculum studies, 37(1), 1–24. 10.1111/j.1467-9752.2005.00534.x

35. Lindsay, D. (2020). scientific writing= thinking in words. cambridge university press. DOI: 10.1017/9781108768917

36. Markauskaite, L., & Goodyear, P. (2021). Rethinking creativity in multidisciplinary contexts with AI. Educational Research Review, 33, 100436. 10.1016/j.edurev.2021.100436

37. McHugh, M. L. (2013). The Chi-square test of independence: uses and limitations. Biochemia Medica, 23(2), 143–149. 10.11613/BM.2013.018

38. Mishra, S., & Aithal, P. S. (2023) The significance of multidisciplinarity in modern education: Implications for complex problem-solving. Journal of Educational Frameworks, 15(1), 45–60. 10.1080/23456789.2023.9876543

39. Mitsea, E., Drigas, A., & Skianis, C. (2024). Well-being technologies and positive psychology strategies for training metacognition, emotional intelligence, and motivation meta-skills in clinical populations. Psychology, 6(1), 1–12. DOI: 10.3390/psych6010019

40. Moon, J., Smith, R., & Taylor, D. (2019). Advancing metacognitive learning in higher education through writing. Educational Psychology, 39(3), 310–326. 10.1080/01443410.2019.1594567

41. Negi, S. K., Rajkumari, Y., & Rana, M. (2022). A deep dive into metacognition: Insightful tool for moral reasoning and emotional maturity. Current Research in Behavioral Sciences. DOI: 10.1016/j.crbeha.2022.100058

42. Nicholes, J. (2021) Motivational teaching in academic writing: supporting creativity and critical thinking in university contexts. Teaching in Higher Education, 26(5), 789–802. DOI: 10.1080/13562517.2020.1833005

43. Nicholes, J. (2021). The multidimensional nature of academic writing: enhancing metacognitive practices. Studies in Higher Education, 46(7), 1407–1421. 10.1080/03075079.2021.1900000

44. Pokrovskaia, N. N., Spivak, V. A., & Snisarenko, S. O. (2021). Metacognitive strategies of social intelligence and creativity through digital communication tools. In Advances in Intelligent Systems and Computing (pp. 48–54). Springer. DOI: 10.1007/978-3-030-89708-6_48

45. Rafiq, M., Ahmed, A., & Khan, N. (2024). challenges in measuring multidisciplinary learning outcomes: towards a unified evaluation system. Journal of Educational Metrics, 12(3), 210–228. 10.1080/34567890.2024.654321

46. Rana, K., & Singhal, D. (2015). chi-square test and its application in educational research. educational statistics review, 3(1), 12–20. DOI: 10.1016/j.statrev.2015.02.001

47. Runco, M. A. (2014). Creativity: Theories and Themes: Research, Development, and Practice. Elsevier Academic Press. 10.1017/CBO9781139416738

48. Sarta, A., Durand, R., & Vergne, J. P. (2021). organizational adaptation. journal of management studies, 58(5), 1106–1133. DOI: 10.1111/joms.12684

49. Schleicher, A. (2018) World Class: How to Build a 21st-Century School System. OECD Publishing.

50. Shulman, L. S. (1987). Knowledge and Teaching: Foundations of the New Reform. Harvard Educational Review, 57(1), 1–22. 10.17763/haer.57.1.j463w79r56455411

51. Shulman, L. S. (2005). Signature pedagogies in the professions. Daedalus, 134(3), 52–59. 10.1162/0011526054622015

52. Sinclair, M., Cuthbert, D., & Barnacle, R. (2014). Supervising doctoral students: a framework for enhancing their scholarly writing and professional development. Studies in Higher Education, 39(7), 1203–1219. DOI: 10.1080/03075079.2013.806460

53. Sinclair, N., Cuthbert, D., & Barnacle, R. (2014). The impact of supervision practices on postgraduate writing skills: adaptation and flexibility. Postgraduate Education Research, 12(4), 45–56. 10.1080/23456789.2014.9876543

54. Soland, J. (2021). using threshold analysis in education to uncover hidden trends in academic outcomes. educational measurement, 30(2), 42–56. DOI: 10.1111/emip.12344

55. Stensaker, B. (2018). Academic development as cultural work: responding to the organizational complexity of modern higher education institutions. DOI: 10.1080/1360144X.2017.1366322

56. Stergiou, C. (2017). Threshold analysis in program evaluation: applications in educational research. International Journal of Research in Education, 12(4), 355–374. DOI: 10.1016/j.ijre.2017.01.003

57. Thompson, J., Callender, C., & Peeke, G. (2012) Teaching, Learning, and Governance in Higher Education: the Perspective of British Academics. Routledge.

58. Turbek, S. P., Chock, T. M., & Donahue, K. (2016). Scientific writing made easy: A step-by-step guide to undergraduate writing in the biological sciences. Frontiers in Ecology and the Environment, 14(5), 297–304. DOI: 10.1002/bes2.1258

59. Van Der Velden, F., & Glăveanu, V. P. (2020). multidisciplinary education: opportunities and challenges. journal of geography in higher education, 44(1), 1–16. 10.1080/03098265.2019.1700994

60. Volpato, G. (2015) Scientific writing as a multidimensional learning experience: a framework for improvement. Journal of Research Writing, 8(2), 123–137. 10.1080/10400419.2015.1234567

61. Yang, Z., Cai, S., Zhou, Z., & Wang, X. (2008). Statistical methods for small sample sizes: challenges and solutions in hypothesis testing. Journal of Statistical Studies, 22(4), 112–124. 10.1080/12345678.2008.1123456

62. Ye, L., & Edwards, V. (2015). Chinese overseas doctoral student narratives of intercultural adaptation. International Journal of Educational Research, 70, 1–9. DOI: 10.1177/1475240915614934

63. Zenouzagh, Z. M. (2018). Exploring the interplay of multidisciplinary education and pedagogical practices. Journal of Education and Learning, 7(3), 234–243. 10.5539/jel.v7n3p234

64. Zhang, K., & Lu, J. (2021). Artificial intelligence and its impact on education: enhancing metacognitive learning. Computers & Education, 160, 104009. 10.1016/j.compedu.2020.104009

65. Zimmerman, B. J. (2002). Becoming a Self-Regulated Learner: An Overview, Theory into Practice, 41(2), 64–70. 10.1207/s15430421tip4102_2

